## Supplemental Information for "A rat epigenetic clock recapitulates phenotypic aging and co-localizes with heterochromatin-associated histone modifications"

**SUPPLENTAL INFORMATION**

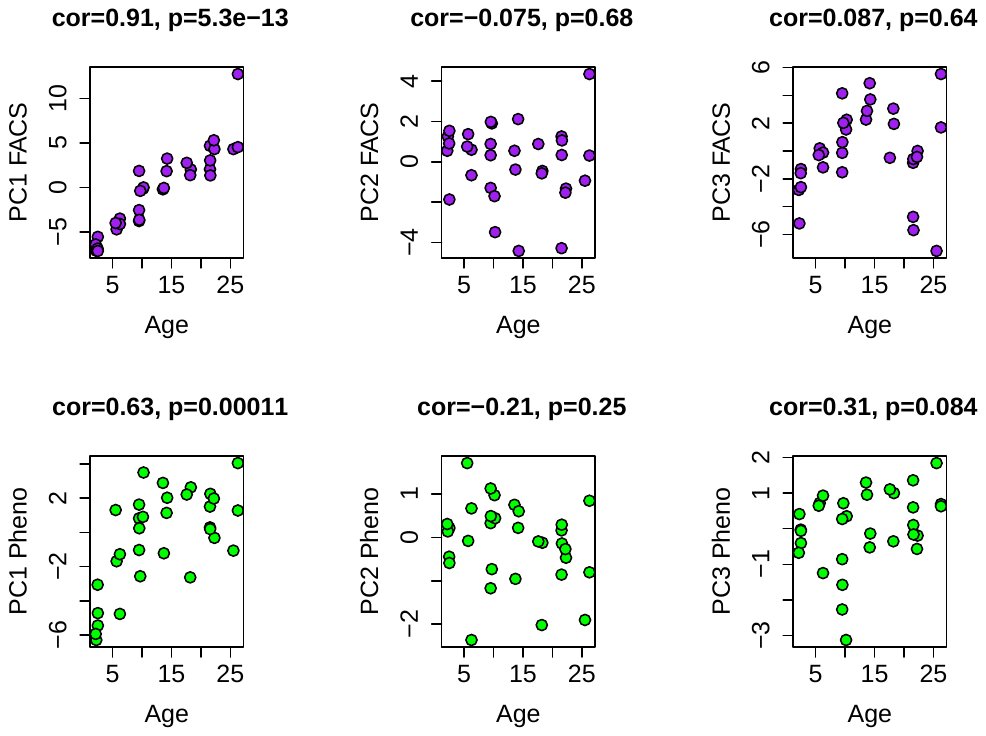

**Fig S1: Age associations for PCs1-3 from FACS and Phenotype data.**

PCs were generated from 39 variables of white blood cell composition using FACS (top row) and from ten phenotypic variables (bottom row), which included eight summarized open field test variables and two rotarod variables (max time and mean time). In both instances, only PC1 is strongly correlated with age.

**Table S1: PC1 Loading for Phenotypic Variables**

| Variable | PC1 Loading |
| --- | --- |
| open field - time ambulatory (s) | 0.3966 |
| open field - horizontal activity count | 0.3872 |
| open field - distance traveled (cm) | 0.3674 |
| open field - vertical activity count | 0.3646 |
| open field - time climbing (s) | 0.3629 |
| rotarod - mean time | 0.2325 |
| rotarod - max time | 0.2299 |
| open field - time in center (s) | 0.1729 |
| open field - time in stereotypy (s) | 0.0569 |
| open field - time resting (s) | -0.3915 |

**Table S2: Phenotypic Associations with DNAmAge based on Overlapped CpGs**

|  | | Beta Coefficient (P-value) | | |
| --- | --- | --- | --- | --- |
|  | | Model 1 | Model 2 | Model 3 |
| DNAmAge (PC1) | |  |  |  |
|  | Age | 0.54 (4.3e-15) | 0.45 (3.9e-12) | 0.29 (2.6e-4) |
|  | Pheno PC1 | -- | 0.43 (4.7e-4) | 0.39 (7.0e-4) |
|  | FACS PC1 |  | -- | 0.31 (1.5e-2) |

**Table S3: Association Between DNAmAge and Caloric Restriction (CR) in C57BL/6**

|  | | Beta Coefficient (P-value) | |
| --- | --- | --- | --- |
|  | | Model 1 | Model 2 |
| DNAmAge~ | |  |  |
|  | Age | 0.16 (<2e-16) | 0.16 (<2e-16) |
|  | CR | -1.21 (1.6e-4) | 1.18 (2.2e-1) |
|  | Age*CR | -- | -0.12 (9.1e-3) |

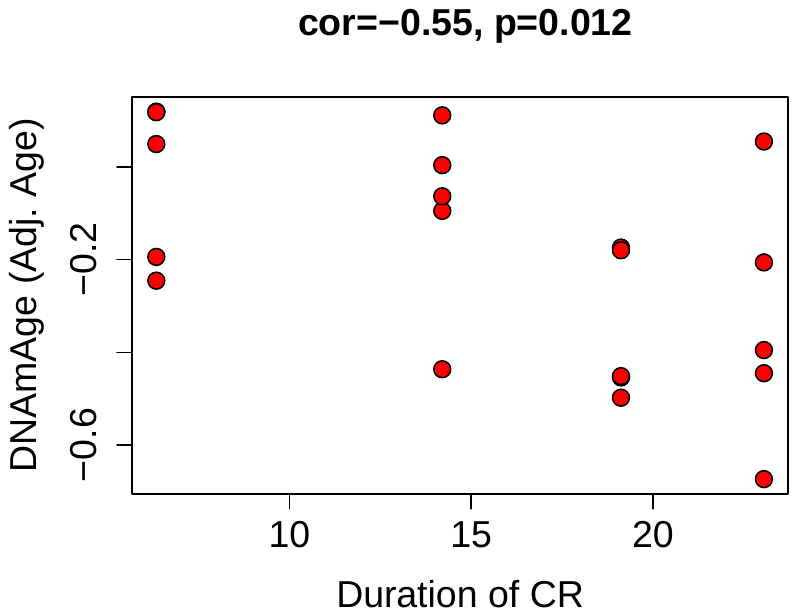

**Fig S2: Correlations between duration of caloric restriction and epigenetic age acceleration.**

Based on the equation of DNAm regressed on age in ad libitum fed mice, we calculated age adjusted DNAmAge for calorically restricted animals. We found that this was correlated with duration the animal had been on CR, suggesting that a longer time spent on the intervention was associated with a greater reduction in epigenetic age.

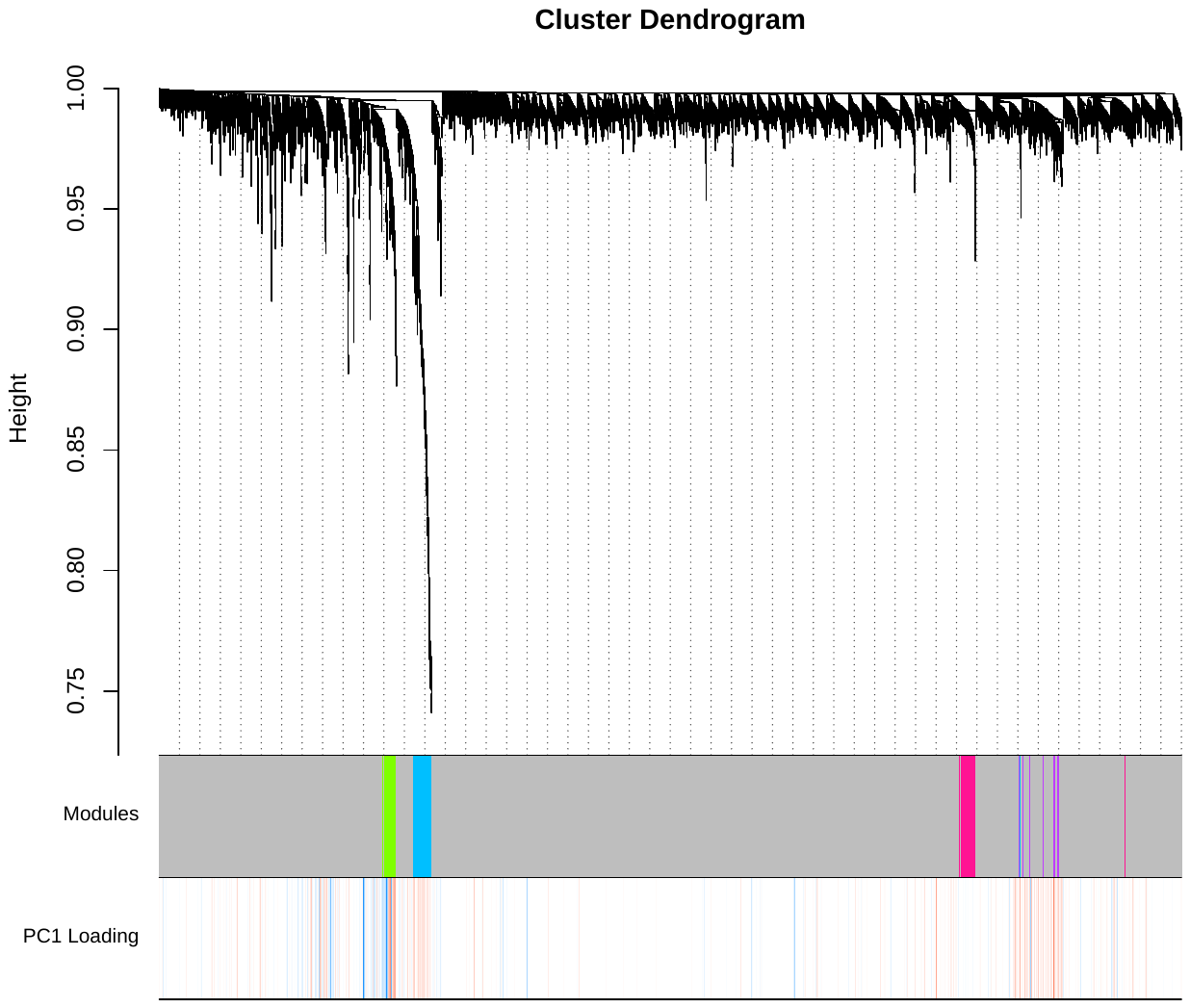

**Fig S3: Dendrogram from WGCNA.**

The y-axis pertains to distance, such that lower values pertain to higher topological overlap. The first column displays the module colors for each CpG—the majority of CpGs are unassigned (grey module), whereas a smaller proportion are in the green, blue, pink, or purple modules. PC1 loadings for the PC used for DNAmAge is shown in the second column (red is positive, blue is negative). CpGs in the purple and blue module tend to have strong positive loadings, whereas green has a mix of strong positive and negative loadings, that segregate based on the dendrogram.

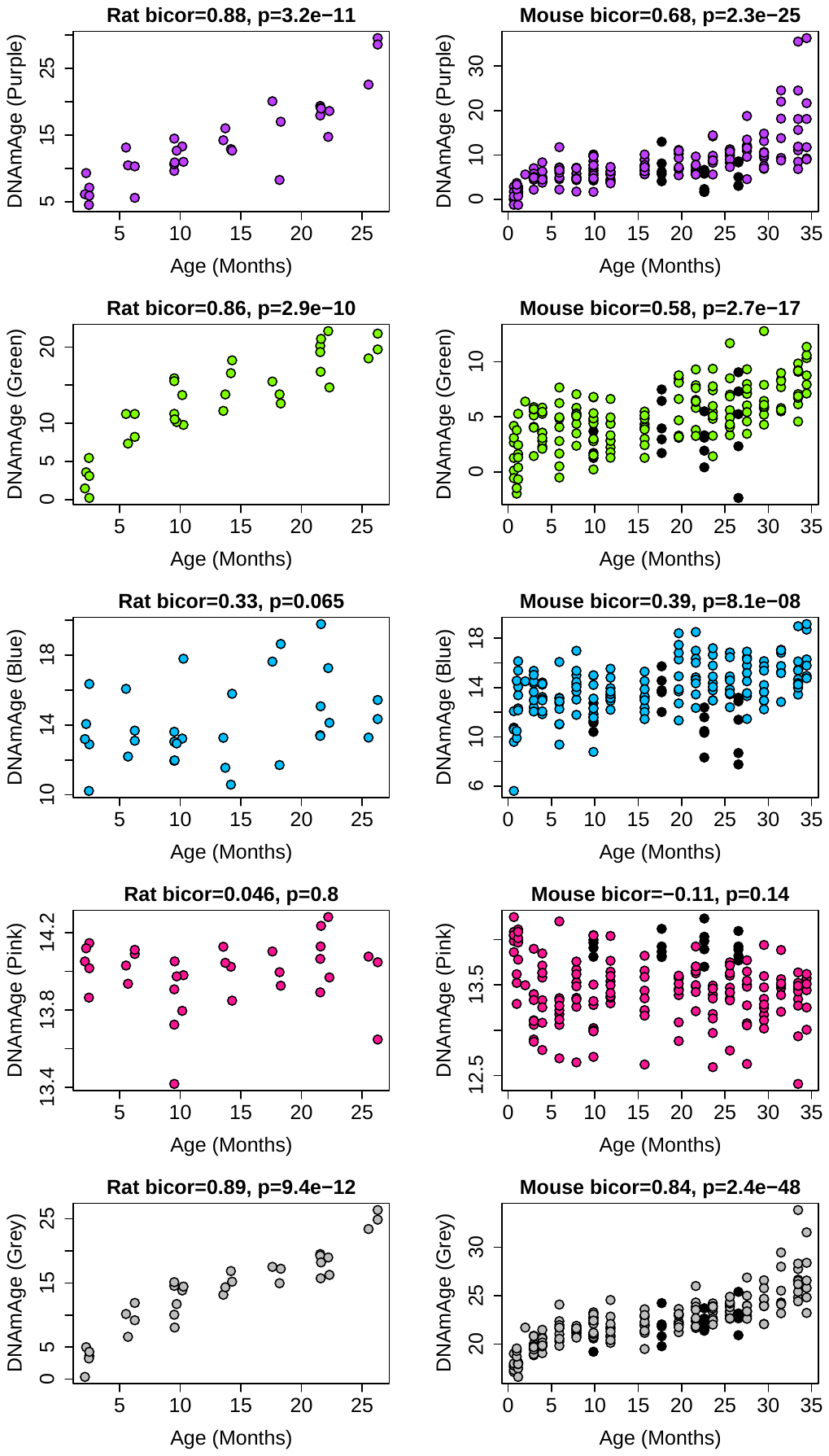

**Fig S4: Correlations between module-based DNAmAge and age in both rats and mouse.**

Modules are denoted by color. Panels in the first column show biweight micorrelations between age and DNAm Age in the validation set for rats. Panels in the second column show biweight micorrelations for C57BL/6 Mice. Black dots in the right panels denote samples who underwent caloric restriction starting at 14 weeks.
